# Use of multimodal sensory cues in predator avoidance by wild-caught zebrafish shoals

**DOI:** 10.1101/2023.01.16.524267

**Authors:** Ishani Mukherjee, Aniket Malakar, Dipjyoti Das, Anuradha Bhat

**Author notes:** Correspondence Address: Department of Biological Sciences, Indian Institute of Science Education and Research (IISER), Kolkata, Mohanpur – 741246, WB, India.

## Abstract

Shoaling in fishes is regulated by factors like predation, vegetation cover, water flow and food availability. Shoals detect and respond to changes in these ecological factors using a multimodal sensory system. Here, we examine the immediate response of wild-caught zebrafish (*Danio rerio)* shoals to cues from its natural predator, the snakehead (*Channa sp*.). Zebrafish shoals were recorded upon exposure to (1) olfactory predator cues, (2) visual predator cues, (3) both cues together, and (4) no cue. We tracked individuals and analysed shoal responses across these treatments. We found that compared to control treatments, shoals receiving either visual or olfactory cues had significantly greater: (i) cohesion, (ii) polarization and (iii) velocity. Interestingly, when the shoals received both cues simultaneously, the cohesion, polarization and velocity decreased and a significantly greater number of individual freezing events occurred. Therefore, zebrafish relied on both visual and olfactory cues to escape predation. However, when shoals were presented with both cues together, while freezing frequency increased, other responses were comparable to control treatments where no predator cue was provided. While this study indicates that multimodal cues elicit a different anti-predator response than the cues singly, more experiments are required to identify the underlying cause of this behaviour.

## Introduction

Group-living provides a variety of benefits to animals, such as, reduced chances of predation, increased foraging efficiency, increased vigilance and faster information transfer (Rubenstein 1978; Krause et al. 2002). However, it also brings disadvantages like the potential increase in the spread of parasites and competition for food or mates (Rubenstein 1978; Rifkin et al. 2012). Evolution of group-living is, thus, a result of the tradeoffs between the costs and benefits associated with grouping. Various ecological factors like predation, habitat structure and resource availability drive group living and impact group level properties (Herbert-Read et al. 2017; Ofstad et al. 2016; Johnson et al. 2002). As a result of recent human-driven changes to their natural habitats, animal groups are often unable to employ optimal predator evasion strategies (Kroeker et al. 2014; Malavasi et al. 2013; Laidre et al. 2008). Research on how animals perceive and evade predators would help in not only improving our understanding of physiology and collective behaviour but also aid in formulating better conservation strategies. This study investigates the importance of predator cues on shoal forming dynamics among fish shoals and differential responses to different types of these cues.

Predation is a key driver of group living amongst many species. In the presence of predators, individuals within groups benefit through dilution effect, group vigilance, confusion effect and selfish herding (Rubenstein 1978; Lehtonen and Jaatinen 2016; Krause et al. 2002). Under predation risk, groups can, along with an increase in group size, developed a suite of other antipredator behaviour (Hass and Valenzuela 2002). Experiments involving juvenile chub (*Leuciscus cephalus*) indicated that predation, unlike competition for food, strongly controls shoal properties such as size-assortativeness (Krause 1994). Predation may, however, also cause individuals to leave groups. In the latter case, this may be explained in terms of an increase in the predator’s search time (for prey) and reduction in prey encounter rates, and thus consequently, lowering of a prey’s chances of being attacked by a predator (DeMars et al. 2016; Hebblewhite and Pletscher 2002).

Prey species may recognize the presence of a predator by perceiving visual, acoustic, olfactory or tactile signals from a predator (Carthey and Blumstein 2018). Further, prey species often rely on multimodal cues to evade predator attacks. Desert tortoises (*Gopherus agassizii*), showed a greater anti-predator response when exposed to visual and olfactory cues from a coyote (*Canis latrans*), as compared to when exposed to only coyote odour (Nafus et at. 2017). Tadpoles of neotropical poison frog (*Allobates femoralis)* can effectively avoid predators only when both visual and olfactory cues from the predators are present (Szabo et at. 2021). Mosquitofish (*Gambusia holbrooki*) detect a predator better when both visual and olfactory cues of a predator are provided (Ward and Mehner 2010).

In freshwater habitats, however, acquiring information on the presence of predators, food, or familiar conspecifics can be challenging for organisms. Visual cues are insufficient if the habitat has frequently changing turbidity levels or is too deep (Utne-Palm 2002) and furthermore, olfactory cues can get diluted in flowing water bodies. This suggests that a combination of olfactory and visual cues would be critical for survival in such habitats. For example, in foraging sticklebacks (*Gasterosteus aculeatus*), olfactory cues can provide the initial information about the presence of prey, following which visual cues provide the fish with a more precise idea of the prey’s location (Johannesen et al. 2012). In such aquatic habitats, prey species also face challenges, where their chances of survival depend on their ability to detect and evade predators. Experiments show that gobies (*Asterropteryx semipunctatus*) and cichlids (*Neolamprologus pulcher*) respond similarly to visual and olfactory cues from a predator (McCormick and Manassa 2008; Fischer et al. 2017). However, in gobies (*Asterropteryx semipunctatus*), the response (in terms of decrease in individual feeding strikes) was greater when both cues are present (McCormick and Manassa 2008).

Shoaling in fish species may be driven by both genetics and environment. Guppy (*Poecilia reticulata*) shoals collected from both high and low predation regimes were found to show increased cohesion when treated with olfactory cues (‘alarm substance’) from a predator. Individuals of the F1 generation from the high predation regime continued to form tight shoals, revealing shoaling also has a strong genetic basis in guppies (Huizinga et al. 2009). Further, guppies from high predation habitats not only showed higher shoal cohesion but also differed in their social behaviour and decision-making abilities compared to those from low predation habitats (Herbert-Read et al. 2017; Ioannou et al. 2017). In zebrafish, while shoal traits like interindividual distances have been found to differ among populations from habitats that differ in predation pressure (Bhat et al. 2015; Suriyampola et al. 2016; Séguret et al. 2016), it is not clear if these are driven by genetic differences or whether these are likely to be more context dependent (i.e., environment-driven/plastic) in nature. Shoal plasticity in response to threat has been studied in some species. Fathead minnow (*Pimephales promelas*) shoals, on detecting threat, voluntarily signal disturbance cues to shoal members to induce tighter shoaling along with behaviours like dashing (rapid movements) and freezing (reduction in activity) (Bairos-Novak et al. 2019). Zebrafish shoals exhibit predator responses like erratic movements, diving, and freezing bouts-the magnitude of such responses depend on the dose of the alarm substance (Speedie and Gerlai 2008). On receiving visual cues from a predator too, zebrafish exhibit similar antipredator behaviours, and observing conspecifics exhibiting such defensive behaviours trigger antipredator behaviour along with increase in cortisol levels within individuals (Oliveira et al. 2017).

Here, we investigated the immediate context-dependent response (plasticity) exhibited by wild-caught zebrafish (*Danio rerio*) shoals, to cues from a natural predator (here, the snakehead, *Channa spp*.). We conducted laboratory-based experiments to examine prey response to only visual, or olfactory and both cues from a snakehead, and measured group level properties such as inter-individual distances and polarization of of wild-caught zebrafish shoals. Analysing video-tracked data on shoal movement, we also examined individual trajectories and measured individual velocity, frequency of freezing events and the proportion time spent in individual freezing events (cessation of movement for a minimum of 5 frames), a typical anti-predator behaviour in zebrafish (Kalueff et al. 2013). Thus, our study focused on collective properties of shoals and freezing behaviour of individuals within these shoals. While olfactory cues provide information on the presence of the predator, visual cues give information regarding position of the predator, and, therefore, we asked whether the separate pieces of information (only olfactory/visual cues) elicit a similar response. We also asked whether shoals show greater antipredator responses when both pieces of information are provided simultaneously.

Earlier studies show that zebrafish use both cues for recognition of familiar individuals, as well as during learning and detecting predators (Spence et al. 2008). In terms of foraging, zebrafish (AB strain) rely primarily on visual cues (Howe et al. 2018), and in turbid environments, visual landmarks improve the foraging efficiency of zebrafish (Sekhar et al. 2019). As zebrafish are known to occur in habitats that vary considerably in turbidity and flow, the magnitude of visual and olfactory information available to the fish varies both temporally and across habitats (Spence et al. 2008). Thus, to survive in the wild, individuals would require strong mechanisms for olfactory as well as a visual perception. As having the ability to process visual and olfactory signals would improve both the quality and magnitude of information available to individuals in the shoal, we hypothesized that zebrafish shoals would depend strongly on olfaction as well as vision to evade a predator. We predicted that shoal responses would be enhanced in the presence of the two cues when available together, as compared to the presence of either of these cues considered separately.

## Methods

### Maintenance of fish

Wild-caught zebrafish from the Ganges drainage in Howrah district (West Bengal, India) were purchased from a local collector. In their natural habitats around West Bengal, these fish typically occur at a temperature range of 23°C-30°C, a pH range of 6.8-8.4, conductivity range of 460μS/cm - 500 μS/cm and a TDS range of 210ppm-265ppm (pers. obs.). Their habitat comprise of moderate vegetation, shallow water with negligible flow. The fish face moderate predation pressure from species like *Channa punctatus, Channa orientalis, Anabas testudineus*. After collection, the fish were transported to the laboratory in aerated plastic bags. In the laboratory, fish were housed in mixed-sex groups of ∼50 adult individuals in 30cm×30cm×60cm bare, aerated glass aquarium tanks, set at 23°C -25°C and a constant lighting condition of 12h dark:12h light in the holding room. The standard body length (tip of snout to base of tail) of the adult individuals were 1.8cm -2.6cm. Fish were fed daily *ad-libitum* with freeze-dried bloodworms or brine shrimp *(Artemia sp*.). To provide olfactory cues from a predator to some shoals, we obtained three similar-sized (body length∼12cm) snakeheads (collected from similar habitats in West Bengal, India) and housed them individually in 20cm×20cm×30cm bare glass tanks. They were maintained under the same laboratory conditions as the zebrafish and were fed with standard pellet food or zebrafish that died naturally (once in a week). To make sure that the predator were not satiated and that (predator) odor cues were consistent over trials, *Channa* individuals were not fed dead zebrafish on the day or on the day before the conduction of experiments.

Shoal responses were recorded under four different predator cue treatments- (1) Controls or no cue treatments (NC) in which shoals received no cues; (2) Olfactory cue treatments (OC) received water from a *Channa* tank. 6.5l of water with predator olfactory cues was added very gently into the center of the arena in these treatments. Often for prey species to elicit anti-predator responses and evade a predator, prior exposure to predator cues is required (Ferrari et al. 2010; Kelley and Magurran 2003; Brown 2003; Mathis and Smith 1993). Wild-caught zebrafish shoals, prior to their collection, have been exposed to odour cues from predators (commonly, *Channa* spp.) in their natural habitats and thus, we expect the shoals to elicit anti-predator responses. It is established that water from a tank housing a predator contains olfactory cues from the predator that evokes antipredator responses in prey (Fischer et al. 2017; Stratmann and Taborsky 2013). Based on this, we added water from a *Channa* tank in treatments receiving olfactory cues from a predator. We calculated (based on prior tests, details in the Supplementary Material) that addition of 6.5l odour cues to the water would simulate the concentration of odour cues a 12cm-sized predator generates in a shallow stagnant pond. Control experiments performed showed that gentle addition of water into the arena center had no impact on shoaling properties (details of experiment and results are in Supplementary Material, Figure S1). (3) Visual cue treatments (VC) in which shoals received visual cues from a toy model predator that resembled a generic predator belonging to the genus *Channa*. Experiments performed showed that test zebrafish respond similarly to the toy predator and a live one and their responses are different when presented with a novel object (details of experiments and their results are in Supplementary Material, Figures S2A, S2B and S2C). The toy predator visually resembled the live predators from which we obtained predator olfactory cues: it was 12cm in length and had horizontal stripes. Thus, this ensured a controlled experimental set up wherein, for OC and VC treatments we exposed all shoals to odour and visual cues of a similar intensity of a perceived threat. (4) Visual and olfactory cue treatments (VOC) in which shoals received both visual and olfactory from a predator. In this case, the toy predator and 6.5l of water containing olfactory cues from the tank housing the *Channa* were added.

All experimental trials were performed between 11:00-15:00 hours every day. A mixed-sex shoal, comprising ten individuals was gently transferred into a 75cm×75cm×12cm arena filled with aged water up to a depth of 4cm or 5cm. Individuals that formed the shoal were chosen randomly from the stock tank and as the stock tank had roughly equal number of males and females (the sex ratio of this particular population was neither male or female skewed), a similar male to female ratio constituted the shoals. It is well known that sex of individuals impact shoal cohesion and shoaling preferences (Snekser et al. 2010; Herbert-Read et al. 2017) and thus, by randomly picking individuals (and maintaining roughly equal number of males and females) to form a shoal, we ensured that the impact of sex of individuals on shoal properties is negligible. The arena was lit by placing two 20W LED light bulbs on either side. The shoal was allowed to acclimatize for 20 minutes following which predator cue(s) were added or not added, depending on the treatment regime. Shoals that would receive olfactory cues from the predator were put in the arena with water depth ∼4cm as the final water depth would become 5cm after addition of olfactory cues. Thus, the total volume of water in the arena was maintained to be constant across the treatments (i.e., control and different predator cue treatments). Two minutes after addition of cues, (to allow shoals to recover from the disturbance caused by addition of cues - if any), or immediately after acclimatization in case of NC, the shoal was video recorded for 5 minutes, using a camera (Canon Legria HF R306) placed vertically overhead. To remove potential effects of residual chemical cues (of conspecifics or of a predator), between consecutive trials, the arena was completely emptied and rinsed with aged water. We tested 60 shoals, 15 per treatment and each shoal was tested once. The order of treatment-type was random and all analyses was performed by the observer blind to the treatment.

### Data preparation and statistical analysis

All data on shoal properties were obtained from scoring of the video recordings of the shoals across treatments. To quantify shoal cohesion, we calculated inter-individual distances between shoal members every 10s. Inter-individual distances between all shoal members were obtained every 10s from coordinates of the entire shoal (following Michael et al. 2021; Kimbel and Morell 2015). The mean inter-individual distance for a given shoal was the mean of inter-individual between all individuals. We measured inter-individual distances every 10s, to ensure an accurate representation of shoal cohesion. In order to visualize cohesiveness among individuals within shoals, we constructed heat plots where the distribution of individuals (in the arena) around a focal individual were plotted (details in Supplementary Material). Next, for each shoal, we calculated the mean polarization score across 30 frames. The polarization score within each frame was measured as the fraction of the number of individuals aligned in the same direction per number of individuals in the shoal (Allan and Pitcher 1986; Miller and Gerlai 2012). Thus, polarization ranges from 0 (no alignment) to 1 (all fishes are aligned). The mean polarization score for each shoal was calculated thereafter by averaging the shoal polarization across 30 frames. We then performed a detailed analysis of the velocity profile of each fish across all shoals. For this, all individuals were tracked using the video-tracking software, ‘idTracker’ (Pérez-Escudero et. al, 2014). The software provided us a tracking accuracy in the range of 70%-99% and thus, using an assisting software, ‘idPlayer’, we manually corrected these videos to improve this accuracy to almost 100%. After this, tracks were smoothed using a moving average of 10 (de Bie et al. 2020) and using MATLAB (R2021a) we plotted the velocity profile of each fish at a temporal resolution of 25 frames per second (details of calculation of velocity has been provided in the Supplementary Material). The mean velocity of the shoal was then calculated using the velocity of each individual. We divided velocity profiles into distinct acceleration and deacceleration phases (based on Harpaz et al. 2017). Our criteria for dividing the velocity into these two phases is described in the Supplementary Material. In addition to these phases, we observed a third distinct phase: freezing, in which individuals were still or moved no greater than half body length (i.e. 1.25cm) in 1s. Freezing behaviour is described similarly by Kalueff et al. 2013. We counted the frequency of such freezing events manually. Next, using MATLAB scripts, we also calculated the proportion time spent by individuals in either of the three states. For comparing the mean shoal velocity across treatments, a VOC shoal was discarded as it had a poor tracking accuracy and hence for this particular result, the sample size was 15 shoals for NC, OC and VC and 14 shoals for VOC. Outlier data (data points outside three standard deviations from the mean (Ratcliff 1993)) for proportion time spent freezing, were discarded from statistical analyses, resulting in sample sizes of 14 shoals for NC, OC, and VC shoals, and a sample size of 13 shoals for VOC treatments respectively. For all other analyses, the sample size was 15 shoals across all 4 treatments.

All statistical analyses were performed using R, version 4.0.2 (R Core Team 2020). We compared shoal cohesiveness across treatments as shown by heatplots, based on Chi square tests (followed by Bonferroni correction to obtain adjusted p values for multiple paired comparisons) for differences in proportional area (of the arena) covered by 25%, 50% and 90% of the shoal across the four treatments. Generalized linear models (GLM) were built to understand the effect of treatment, i.e., cue provided on shoal properties. In each model, a collective shoal trait (mean inter-individual distance, or mean polarization score), the frequency of individual freezing events or the proportion time spent freezing was the dependent variable, and treatment (cue) was the fixed factor. Before constructing all models, we checked the distribution of our data using the ‘fitdistr’ function (Delignette et al. 2005). Model comparisons were performed using ANOVA in ‘car’ package (Fox and Weisberg 2019) and post hoc paired tests (Tukey’s post hoc HSD Test) were carried out for comparing the effects of factors that were significant. Two tailed p-values less than or equal to 0.05 were considered significantly different.

## Results

### Shoal cohesion and polarisation

Shoal cohesion, examined through mean inter-individual distances and heat plots indicated that shoals receiving predator cue(s) were more cohesive compared to shoals in control treatments that did not receive predator cues. Interindividual distances in shoals receiving any form of predator cue(s) i.e., OC (mean±SE, 22.7±1.49cm), VC (mean±SE, 21.85±1.34cm) and VOC (mean±SE, 25.46±0.97cm), was significantly lower compared to NC shoals (mean±SE, 33.13±1.41) (GLM: Wald type IIχ^2^ = 42.73, df = 3, p<0.001, Table 1a) (Tukey’s Test Results: NC vs. OC: Z-value -5.42, p<0.001, NC vs. VC: Z-value -5.86, p<0.001; NC vs. VOC: Z-value - 3.98, p<0.001; OC vs. VC: Z-value -0.44, p=0.97; OC vs. VOC: Z-value: 1.43, p=0.47; VC vs. VOC: Z-value 1.87, p=0.23) (Figure 1a). The shoal cohesion was visualized in the form of heat plots showing distribution of all neighbours relative to the focal individual. Although statistically not significant (χ^2^= 0.002, df = 6, p = 0.15), as indicated by contour lines, shoals were generally tighter (i.e., more number of individuals occupy a smaller area) in treatments receiving predator cue(s) than in the control treatment and shoals receiving both cues (VOC) appear less tight as compared to shoals receiving a single cue (VC or OC) (Figure 1b). There was a significant effect of treatment on polarization in shoals (GLM: Wald type II χ^2^ = 14.64, df = 3, p<0.01, Table 1b), and shoals that received a single predator cue, OC (mean±SE, 0.62±0.03) or VC (mean±SE, 0.62±0.02), were significantly more polarized than NC (mean±SE, 0.49±0.02). Polarization of VOC (mean±SE, 0.58±0.02) shoals was intermediate and comparable to that of the other treatments (Tukey’s Test Results: NC vs. OC: Z-value = 3.26, p<0.01; NC vs. VC: Z-value = 3.35, p <0.01; NC vs. VOC: Z-value = 2.34, p = 0.08; OC vs. VC: Z-value = 0.09, p= 0.99; OC vs. VOC: Z-value = -0.92, p=0.79; VC vs. VOC: Z-value = -1.01, p =0.74) (Figure 1c).

**Table 1a.** Results of the generalized linear model (GLM) for predicting effect of cue on inter-individual distance.

|  | Estimate | Std. Error | t value | Pr(>\|t\|) |
| --- | --- | --- | --- | --- |
| (Intercept) | 33.12 | 1.36 | 24.36 | <0.0001 |
| OC | -10.42 | 1.92 | -5.42 | <0.0001 |
| VC | -11.28 | 1.92 | -5.86 | <0.0001 |
| VOC | -7.66 | 1.92 | -3.98 | <0.001 |

**Table 1b.** Results of the generalized linear model (GLM) for predicting effect of cue on mean polarization score.

|  | Estimate | Std. Error | t value | Pr(>\|t\|) |
| --- | --- | --- | --- | --- |
| (Intercept) | 0.49 | 0.02 | 17.14 | <0.0001 |
| OC | 0.13 | 0.04 | 3.26 | <0.01 |
| VC | 0.13 | 0.04 | 3.35 | <0.01 |
| VOC | 0.09 | 0.04 | 2.34 | <0.01 |

**Figure 1:**
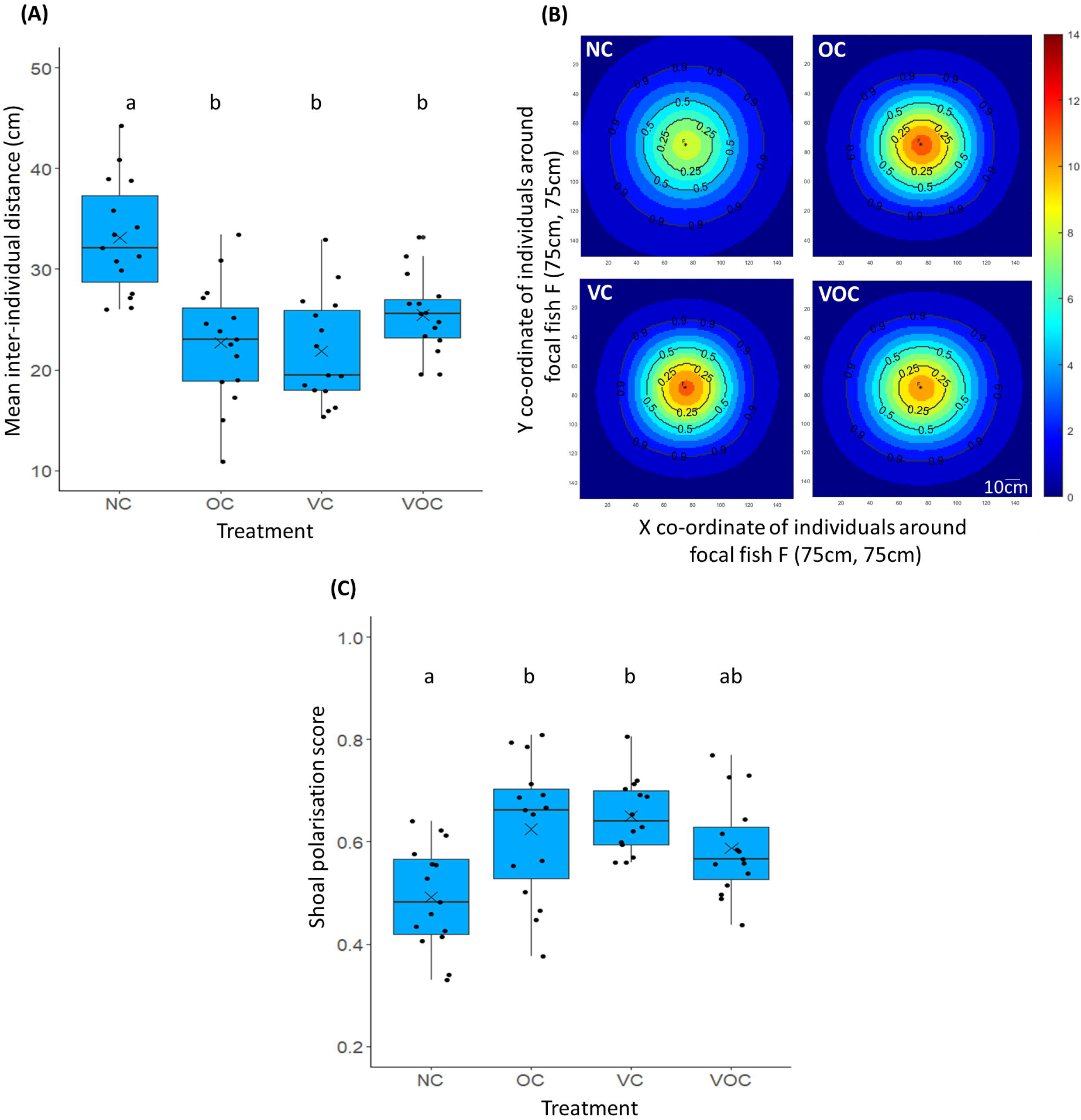
Mean-interindividual distances, distribution of individuals around a focal fish and mean shoal polarization score. **(A)** Box-and-whisker plots across treatments representing the mean interindividual distance for shoals across treatments. **(B)** Heat plots showing the position of individuals around a focal individual positioned at F (75,75). For each treatment, all individuals across all frames have been considered as the focal fish F and the positions of all shoal mates with respect to it are displayed in the form of a density map. The contour lines indicate the proportion of the shoal within a given area. **(C)** Box-and-whisker plots across treatments representing the mean shoal polarization score. In the box and whisker plots, each box represents the interquartile range, the line within the box indicates the median while the whiskers represent the range of the data. Each data point is represented as a dot. The mean is represented as a cross. The different letters placed above the boxes represent significant differences between the categories. Comparisons were performed using Tukey’s HSD Test (N_NC_, N_OC,_ N_VC,_ N_VOC_ =15, p>0.05)

### Shoal velocity

Velocity profile of individuals were subcategorized into acceleration, deacceleration and freezing phases and plotted (for representation, Figure 2a shows the velocity profile of a single individual). Thereafter, we calculated mean shoal velocity and found that it differed significantly across treatments (GLM: Wald type II χ^2^ = 25.51, df = 3, p<0.0001, Table 2a). OC (mean±SE, 6.56±0.37cm/s). VC (mean±SE, 5.98±0.37cm/s) shoals had a significantly greater mean shoal velocity than NC shoals (mean±SE,4.30±0.27cm/s). The mean shoal velocity of VOC shoals (mean±SE,4.78±0.30cm/s) was comparable to NC shoals (mean±SE,4.30±0.27cm/s) and VC shoals (mean±SE, 5.98±0.37cm/s) but significantly lower than OC shoals (mean±SE, 6.56±0.37cm/s) (Tukey’s Test Results: NC vs. OC: Z-value = 4.49, p<0.001; NC vs. VC: Z-value = 3.34, p <0.01; NC vs. VOC: Z-value = 0.91, p = 0.79; VC vs. OC: Z-value = -1.14, p = 0.66; OC vs. VOC: Z-value = -3.41, p<0.01; VC vs VOC: Z-value = -2.30, p=0.09) (Figure 2b).

**Table 2a.**
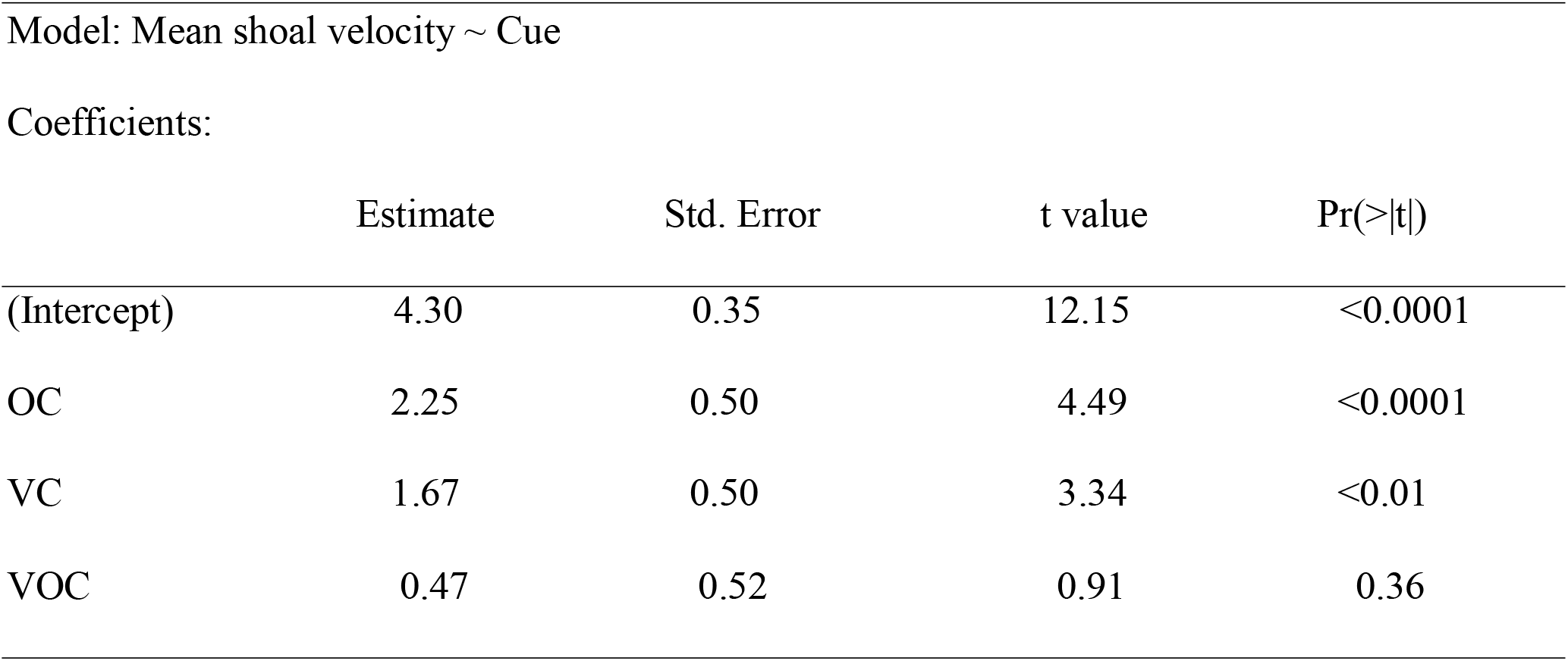
Results of the generalized linear model (GLM) for predicting effect of cue on mean shoal velocity.

**Figure 2:**
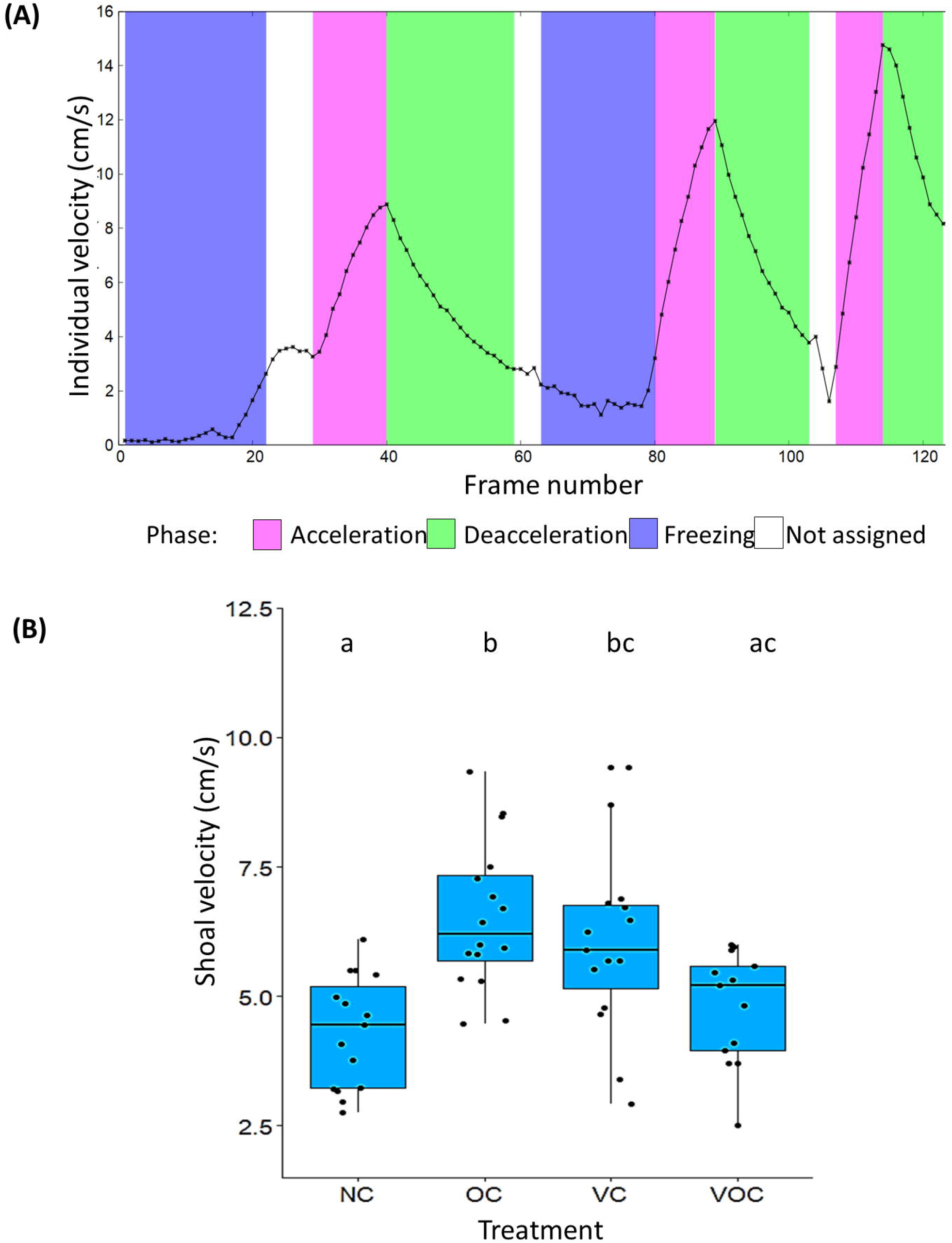
Velocity profile of a single fish, and mean shoal velocity. **(A)** The velocity profile of an individual fish across 10 seconds. Velocity(cm/s) was calculated based on the individual’s trajectory every frame (i.e. every 0.04s). The velocity has been divided into acceleration (shaded pink), deacceleration (shaded green) and freezing phases (shaded indigo). We categorized velocity into acceleration, deacceleration or freezing only when these events continued at least for 5 successive frames. The white gaps indicate “noise” -events that did not last for at least 5 frames. Box-and-whisker plots across treatments representing **(B)** The mean shoal velocity (in cm/s). Each box represents the interquartile range, the line within the box indicates the median while the whiskers represent the range of the data. Each data point is represented as a dot. The mean is represented as a cross. The different letters placed above the boxes represent significant differences between the categories. Comparisons were performed using Tukey’s HSD Test (N_NC_, N_OC,_ N_VC,_ =15, N_VOC_=14; p>0.05)

### Freezing behaviour: frequency of individual freezing events and proportion time spent freezing

Shoals that received both cues showed significantly more freezing frequency than in any of the other treatments (GLM: Wald type II χ^2^ = 25.81, df = 3, p<0.001, Table 3a). VOC (mean±SE, 13.86±2.69) shoals showed significantly more freezing than NC (mean±SE, 4.13±0.55), OC (mean±SE, 3.93±1.38), and VC (mean±SE, 4.86±0.70) shoals (Tukey’s Test Results: VOC vs. NC: Z-value = 4.21, p<0.001; VOC vs. OC: Z-value = 4.29, p <0.001, VOC vs. VC: Z-value = 3.89, p<0.001) (Figure 3a). While there was no significant difference in the proportion time spent in accelerating or deaccelerating (see Supplementary Material for detailed results), GLM revealed proportion of time spent freezing was dependent on the cue provided (GLM: Wald type II χ^2^ = 10.31, df = 3, p=0.01, Table 3b). Post hoc tests, however, showed no significant impact of cues on proportion of time spent swimming. In general, the proportion of time spent freezing in VOC shoals (mean±SE,0.06±0.02) and NC (mean±SE,0.06±0.01) shoals was higher than the proportion of time spent freezing in OC shoals (mean±SE,0.01±0.002) and VC shoals (mean±SE,0.01±0.004) (Figure 3b).

**Table 3a.**
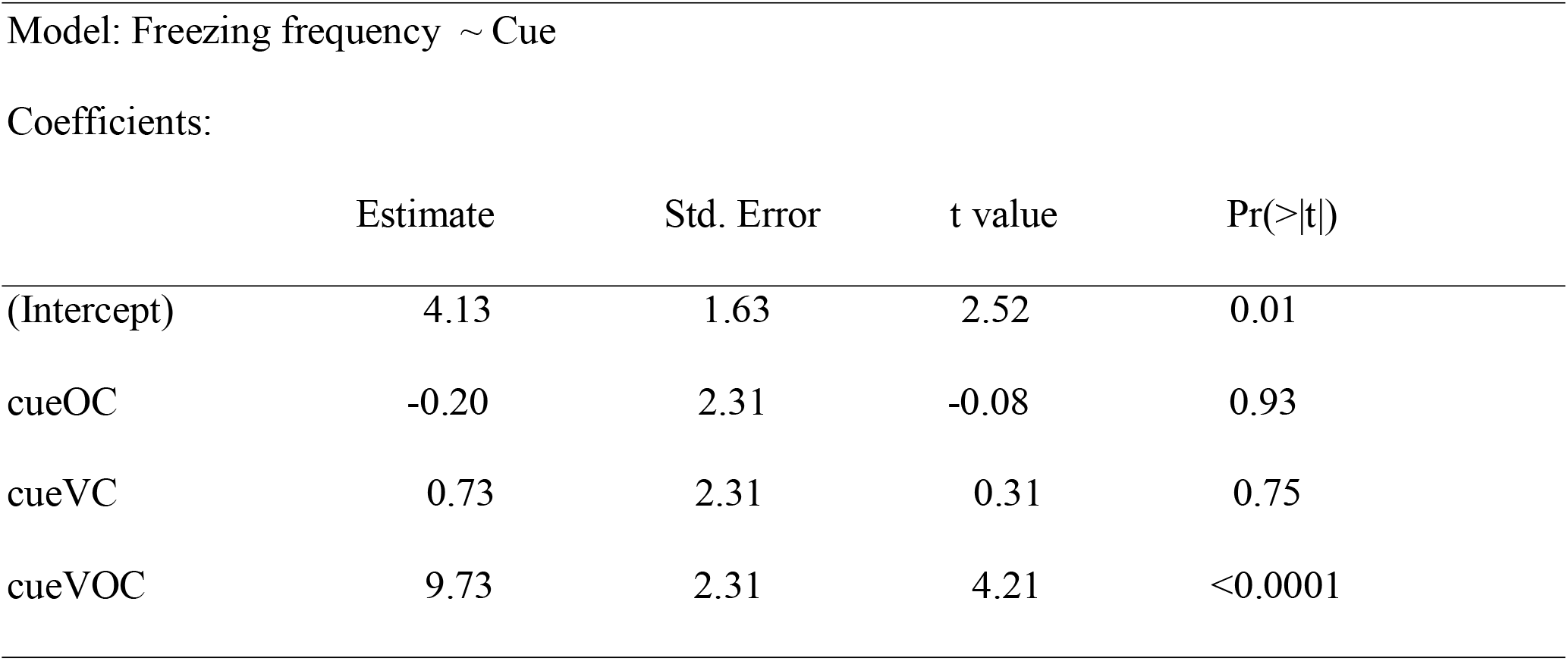
Results of the generalized linear model (GLM) for predicting effect of cue on freezing frequency.

**Table 3b.** Results of the generalized linear model (GLM) for predicting effect of cue on proportion time spent in freezing.

|  | Estimate | Std. Error | t value | Pr(>\|t\|) |
| --- | --- | --- | --- | --- |
| (Intercept) | 0.06 | 0.01 | 3.94 | <0.001 |
| OC | -0.05 | -0.02 | -2.42 | 0.01 |
| VC | -0.04 | -0.02 | -2.27 | 0.02 |
| VOC | -0.002 | -0.02 | -0.12 | 0.90 |

**Figure 3:**
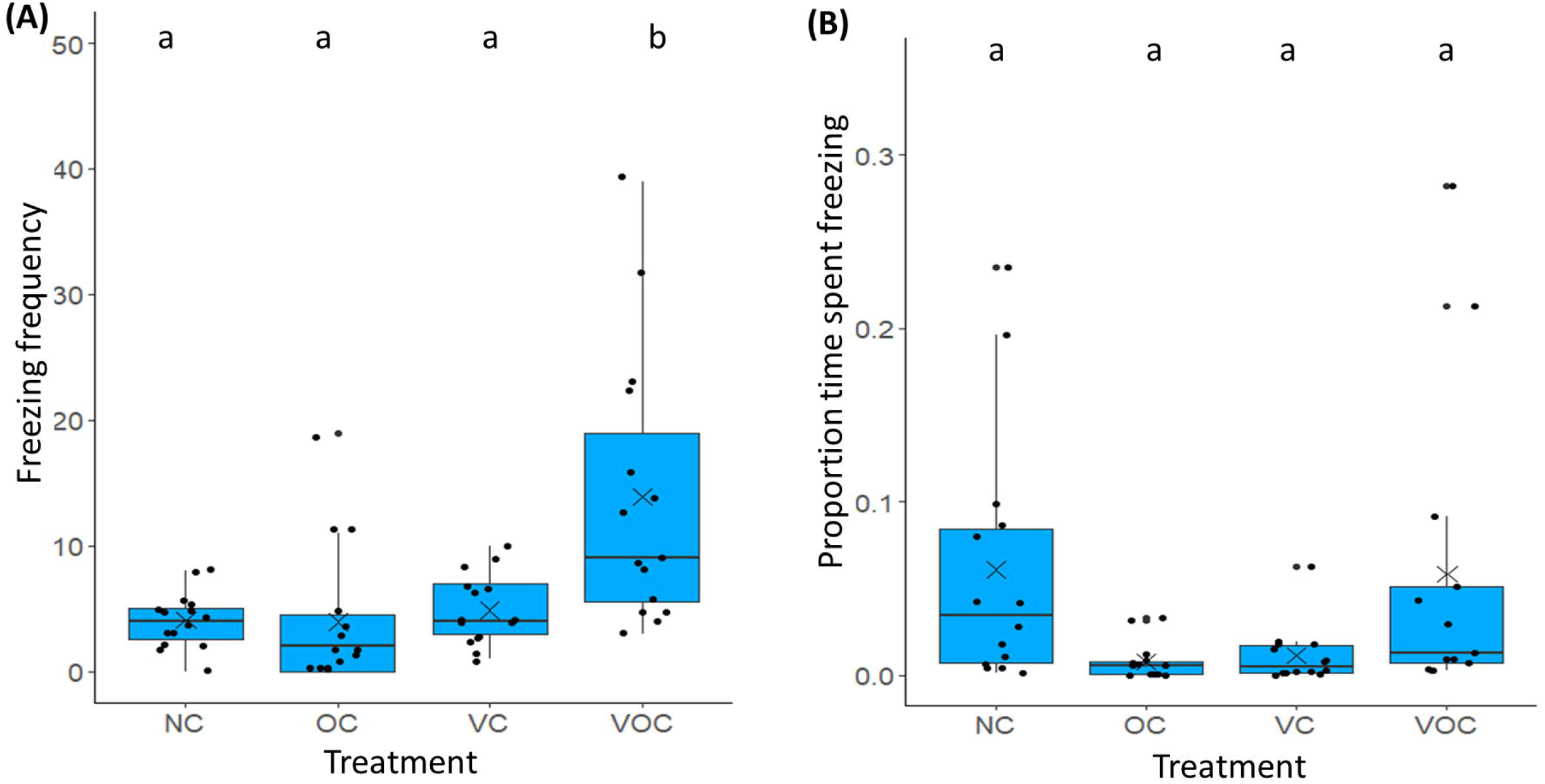
Freezing behaviour: Box-and-whisker plots across treatments representing (A) the frequency of freezing events and (B) the proportion of time spent freezing. Each data point is represented as a dot. The mean is represented as a cross. The different letters placed above the boxes represent significant differences between the categories. Comparisons were performed using Tukey’s HSD Test (For freezing frequency: N_NC_, N_OC,_ N_VC,_ N_VOC_ =15; For proportion of time spent freezing: N_NC_, N_OC,_ N_VC_ =14 ; N_VOC_=13; p>0.05).

## Discussion

Our study revealed that shoal dynamics and their properties in wild-caught zebrafish shoals change considerably as an immediate response to predator cue(s). Interestingly, when exposed to dual predator cues, these shoals increased frequency of freezing and shoal properties of these shoals resembled that of shoals receiving no cues. While this work provides a new insight into predation evasion strategies among wild zebrafish shoals, more experiments are required to understand the benefits of such behaviour and the underlying mechanisms that generate the observed response among these shoals.

In the presence of a single predator cue, shoals exhibited considerable plasticity in the form of increased cohesion, polarization and velocity. As reported for other species like gobies and cichlids (McCormick and Manassa 2008; Fischer et al. 2017), their reliance on either of the visual or olfactory cues for predator evasion is comparable. It must be pointed out that while these responses might be comparable between shoals receiving visual cues and shoals receiving olfactory cues from a predator, other kinds of responses (such as habitat use, predator inspections) might differ. Experiments where anti-predator response was measured in terms of habitat use in roach (*Rutilus rutilus*) revealed that the fish species chose a different kind of habitat based on the type of cue provided: while the fish entered open water habitats in response to odour cues from a predator, the species when exposed to visual cues from the same predator entered a structured habitat (Martin et al. 2010). Gambusia (*Gambusia affinis*), during risky behaviours such as predator inspection, relied solely on visual cues from a predator (the presence of olfactory cues made no difference) (Smith and Belk 2001). In our study, we specifically focused on the dynamics of shoal properties such as cohesion, polarization and velocity in response to either visual and olfactory cues from a predator, and in these properties, we found comparable responses to olfactory and visual cues from a predator.

Interestingly, when shoals were presented with both cues individuals underwent increased freezing and some shoal properties (i.e., polarisation and shoal velocity) were comparable to that of NC shoals. This could be due to the possible effect of incomplete or diffused information transfer within the shoal. In their natural habitat, due to turbidity and a cocktail of olfactory cues, such shoals almost always have incomplete information of a potential predator. To gather information on risk imposed by a predator, prey species integrate reliable as well as unreliable information of the predator (Feyten et al. 2019). Across taxa, prey species show greater fear responses on encountering incomplete information (Ferrari et al. 2012; Crane and Ferrari 2017). Therefore, we speculate that unable to sense the magnitude of threat present, it is possible that these shoals elicit the highest anti-predator response. It is possible that such responses could be beneficial for these shoals as there is a possibility that the level of the threat is the highest. When the shoals have a more clear information about the predator (position in arena and smell) they are less wary of it as they are better informed about a potential threat. Another possible explanation is that in the presence of dual cues, these shoals change their predator avoidance strategy by showing increased freezing, rather than adhering to safety in numbers. We speculate that increased freezing may be a more beneficial strategy to avoid a predator. A third, related possibility is that the shoals are unable to execute the antipredator response as combined cues increases individual stress levels. Thus, when predator cues are highest, the anti-predator response in these shoals fail and shoals might be unable to escape predation, or might resort to freezing. Future experiments to investigate which strategy (safety in numbers or increased freezing) is more benefitial to avoid a ambush predators such as *Channa* might provide insights into which of the above hypotheses holds true. It is well known that cortisol levels in zebrafish increase in response to a predator (Barcellos et al. 2007; Oliveira et al., 2013). Further insights may be obtained by analyzing cortisol levels among these shoals: cortisol levels in shoals receiving dual cues will indicate the stress levels among these shoals.

On receiving cue(s) from a predator, shoal cohesion, polarization and velocity increased. Our results on increased cohesion on exposure to predator cue(s) are in consensus with studies on different fish species (Herbert-Read et al. 2017; Meuthen et al. 2016; Cattano et al. 2019). In recent studies, shoal cohesion in wild zebrafish have been found to be population dependent (Suriyampola et al. 2016; Bhat et al., 2015) and thus, we speculate that in natural habitats, populations with greater predation pressure (than what was presented in the current study) might have even lower inter-individual distances. Observations on polarization in zebrafish shoals indicate a bimodal distribution, which means shoals have two distinct modes of collective motion (Miller and Gerlai 2012). When the polarization is high, fish are said to be schooling and when the polarization is low fish are said to be shoaling. We observed that shoals which received a single predator cue were more polarized. Changing to a more polarized state (schooling) in response to a predator threat has been previously reported in shoals (Herbert-Read et al. 2017; Rieucau et al. 2015; Magurran and Seghar 1990; Magurran and Pitcher 1987). Shoals of zebrafish showed higher polarization when they were introduced to a novel tank, and polarization decreased over time, as shoals got habituated (Miller and Gerlai 2012). This strongly suggests that alignment in the same direction i.e., assuming a polarized state, is an immediate response to perceived threat among zebrafish shoals. In addition to increased cohesion and polarization, like other species such as guppies, the mean velocity of a shoal also increased (Ghalambor et al. 2004; Plaut 2001). Higher velocity in these shoals reflects erratic movements which is a typical antipredator response in zebrafish (Kalueff et al. 2013; Speedie and Gerlai 2008). As a next step, quantification of erratic movements among these shoals (by calculating turn angles) might provide insight into how this particular behaviour changes in reponse to single or dual cues from a predator.

Besides increased cohesion, polarization and velocity, freezing is another strategy often adopted by prey, to avoid a predator. A variety of fish species, including zebrafish, are known to exhibit freezing behaviour in presence of a predator (Seigel et al. 2022, Kimbell and Morell 2015; Kalueff et al. 2013; Malavasi et al. 2013; Brown and Magnavacca 2003, Brown and Dreier 2002). Further, freezing has been shown to increase proportionally with increase in dosage of alarm substance, i.e, predator cue (Speedie and Gerlai 2008). In our experiments, shoals resorted to greater instances of freezing when dual cues were provided to wild-caught zebrafish shoals. The time spent freezing did not differ significanty across treatments indicating that zebrafish shoals exposed to dual cues underwent many short freezing events. Although statistically not significant we observed greater proportion of freezing time in NC and VOC shoals. We speculate that high freezing time when shoals were provided no cues occurs as individuals have been introduced in a novel arena and even after a twenty-minute acclimatization, individuals show some signs of stress in the form of freezing. Controlled experiments with an even longer acclimatization phase may enable us to disentangle the freezing caused by predation from that caused due to fish being exposed to a novel arena. While the longest freezing event we observed lasted for 3.5s (88 frames) in a shoal receiving visual and olfactory cues from the predator, short freezing events that lasted for 0.2s (5frames) were observed among multiple shoals in VC and OC treatments.

Increased freezing by individuals in shoals exposed to dual cues resulted in a temporary decrease in shoal cohesion, polarization and velocity. In consensus with our findings, a previous study on zebrafish shoals exposed to a moving robot fish (considered as a predator) has shown decreased cohesion and increased instances of freezing (Butail et al. 2013). Our study is in consensus to previous studies on zebrafish that clearly show how individual behaviour control physical properties in fish groups.

Besides freezing phases, we also identified acceleration and deacclereation phases among these shoals (similar to Harpaz et al. 2017). We found that mean duration of these phases in wild-caught zebrafish (mean acceleration time= 326ms; mean de-acceleration time= 319ms) are similar to that of lab-bred zebrafish (mean acceleration time= 200ms; mean de-acceleration time= 250ms). A variety of fish species including guppies (*Poecilia reticulata*), dace (*Leuciscus leuciscus*), rainbow trout (*Oncorhynchus mykiss*) and goldfish (*Carassius auratus*) show such acceleration and deacceleration phases-a characteristic of saltatory swimming behaviour (Herbert-Read et al. 2017; Videler and Wardle 1991).

As a natural predator, *Channa* spp. are ambush predators, and usually hide in the muddy substratum and attack a potential prey when it approaches the later (Cagauan 2007; Phen et al. 2004). Although *Channa sp*. are found in the bottom of muddy waters (Amilhat and Lorenzen 2005) and show little movement before striking at the prey, we speculate that some movement (such as steady flapping of pectoral fins) might have alerted shoals more and this is a potential limitation in our experiment.

In the present study, fish used the two cues as separate pieces of information and modified their behavioural response accordingly. Our study shows that individual freezing behaviour impacts group properties such as polarization and velocity. Unlike most studies involving shoal responses to a predator that either report changes in shoal physical properties or behavioural changes, our analyses address both aspects. While this work clearly shows that shoals respond differently to single predator cues as compared to dual predator cues, further studies are needed to understand the benefits of exhibiting such behaviour and why such differences in anti-predator behaviour in response to different cues arise among organisms.

## Author Contributions

A.B. and I.M. conceptualized and designed the experiments; I.M. performed video tracking of fish shoals, analysed the data and wrote the manuscript. A.M. carried out part of the analysis. D.D. gave valuable inputs and helped A.M. in interpreting the analysis. A.B. helped I.M. in performing the analysis, edited the manuscript and supervised the work.

## Acknowledgements

The authors thank the Indian Institute of Science Education and Research Kolkata (IISER Kolkata), India, for providing infrastructural and financial support. The authors thank Mr.Sukanta Bhattacharya for collecting the fishes and Mr. Prasenjit Pan for help in fish maintenance in the laboratory.

## Data availability

The authors are willing to share data associated with this study on reasonable request.

## References

Allan JR, Pitcher TJ, 1986. Species segregation during predator evasion in cyprinid fish shoals. Freshwater Biol 16 (5): 653–659.

Amilhat E, Lorenzen K, 2005. Habitat use, migration pattern and population dynamics of chevron snakehead Channa striata in a rainfed rice farming landscape J Fish Biol 67: 23–34.

BairosLNovak KR, Ferrari MC, Chivers DP, 2019. A novel alarm signal in aquatic prey: Familiar minnows coordinate group defenses against predators through chemical disturbance cues. J Anim Ecol 88: 1281–1290.

Barcellos LJ, Ritter F, Kreutz LC, Quevedo RM, da Silva LB, Bedin AC, Finco J, Cericato L, 2007. Whole-body cortisol increases after direct and visual contact with a predator in zebrafish, Danio rerio. Aquaculture 272(1-4): 774–8.

Bhat A, Greulich MM, Martins EP, 2015. Behavioral plasticity in response to environmental manipu-lation among zebrafish (Danio rerio) populations. PLoS One 10: e0125097.

Borner KK, Krause S, Mehner T, Uusi-Heikkilä S, Ramnarine IW, Krause J, 2015. Turbidity affects social dynamics in Trinidadian guppies. Behav Ecol Sociobiol 69: 645–651.

Brown GE, 2003. Learning about danger: chemical alarm cues and local risk assessment in prey fishes. Fish Fisheries 4: 227–234.

Brown GE, Magnavacca G, 2003. Predator inspection behaviour in a characin fish: an interaction between chemical and visual information?. Ethol 109: 739–750.

Brown GE, Dreier VM, 2002. Predator inspection behaviour and attack cone avoidance in a characin fish: the effects of predator diet and prey experience. Anim Behav 63: 1175–81.

Butail S, Bartolini T, Porfiri M, 2013. Collective response of zebrafish shoals to a free-swimming robotic fish. PLoS One 8(10): e76123.

Cagauan AG, 2007. Exotic aquatic species introduction in the Philippines for aquaculture-A threat to biodiversity or a boon to the economy. J Environ Sci Manag 10: 48–62.

Carthey AJ, Blumstein DT, 2018. Predicting predator recognition in a changing world. Trends Ecol Evol 33: 106–115.

Cattano C, Fine M, Quattrocchi F, Holzman R, Milazzo M, 2019. Behavioural responses of fish groups exposed to a predatory threat under elevated CO2. Mar Environ Res 147:179–184.

Crane AL, Ferrari MC, 2017. Patterns of predator neophobia: a meta-analytic review. Proc Biol Soc B 284 (1861): 20170583.

De Bie J, Manes C, Kemp PS, 2020. Collective behaviour of fish in the presence and absence of flow. Anim Behav 167:151–159.

Delignette-Muller ML, Dutang C, 2015. fitdistrplus: An R Package for Fitting Distributions. J Stat Softw, 64: 1–34.

DeMars CA, Breed GA, Potts JR, Boutin S, 2016. Spatial patterning of prey at reproduction to reduce predation risk: what drives dispersion from groups? Am Nat 187: 678–687.

Ferrari MC, Wisenden BD, Chivers DP, 2010. Chemical ecology of predator–prey interactions in aquatic ecosystems: a review and prospectus. Can J Zool 88 (7): 698–724.

Ferrari MC, Vrtělová J, Brown GE, Chivers DP, 2012. Understanding the role of uncertainty on learning and retention of predator information. Anim Cogn 15(5): 807–813.

Feyten LE, Demers EE, Ramnarine IW, Brown GE, 2019. Predation risk assessment based on uncertain information: interacting effects of known and unknown cues. Curr Zool 65(1): 75–76.

Fischer S, Oberhummer E, Cunha-Saraiva F, Gerber N,Taborsky B, 2017. Smell or vision? The use of different sensory modalities in predator discrimination. Behav Ecol Sociobiol 71: 143.

Fox J, Weisberg S 2019. An R Companion to Applied Regression, Third edition. Sage, Thousand Oaks CA.

Gthalambor CK, Reznick DN, Walker JA, 2004. Constraints on adaptive evolution: the functional trade-off between reproduction and fast-start swimming performance in the Trinidadian guppy (Poecilia reticulata). Am Nat 164(1): 38–50.

Harpaz R, Tkačik G, Schneidman E, 2017. Discrete modes of social information processing pre-dict individual behavior of fish in a group. Proc Nat Acad Sci 114: 10149–10154.

Hasenjager MJ, Hoppitt W, Dugatkin LA, 2020. Personality composition determines social learning pathways within shoaling fish. Proc R Soc B 287: 20201871.

Hass CC, Valenzuela D, 2002. Anti-predator benefits of group living in white-nosed coatis (Nasua narica). Behav Ecol Sociobiol 51: 570–578.

Hebblewhite M. Pletscher, DH, 2002. Effects of elk group size on predation by wolves. Can J Zool 80: 800–809.

Herbert-Read JE, Rosén E, Szorkovszky A, Ioannou CC, et al., 2017. How predation shapes the social interaction rules of shoaling fish. Proc R Soc B 284: 1126.

Herbert-Read JE., Krause S, Morrell LJ, Schaerf TM, Krause J, Ward AJW, 2013. The role of individuality in collective group movement. Proc R Soc B 280: 20122564.

Howe HB, McIntyre PB, Wolman MA, 2018. Adult Zebrafish primarily use vision to guide piscivorous foraging behavior. Behav Processes 157: 230–237.

Huizinga M, Ghalambor CK, Reznick DN, 2009. The genetic and environmental basis of adaptive differences in shoaling behaviour among populations of Trinidadian guppies, Poecilia reticulata. J Evol Biol 22: 1860–1866.

Ioannou CC, Ramnarine IW, Torney CJ, 2017. High-predation habitats affect the social dynamics of collective exploration in a shoaling fish. Sci Adv 3: e1602682.

Johannesen A, Dunn AM, Morrell LJ, 2012. Olfactory cue use by three-spined sticklebacks foraging in turbid water: prey detection or prey location? Anim Behav 84: 151–158.

Johnson DD, Kays R, Blackwell PG, Macdonald DW, 2002. Does the resource dispersion hypothesis explain group living? Trends Ecol Evol 17: 563–570.

Jolles JW, Boogert NJ, Sridhar VH, Couzin ID, Manica A, 2017. Consistent individual differences drive collective behavior and group functioning of schooling fish. Curr Biol 27: 2862–2868.

Kalueff AV, Gebhardt M, Stewart AM, Cachat JM, Brimmer M et al., 2013. Towards a comprehensive catalog of zebrafish behavior 1.0 and beyond. Zebrafish 10: 70–86.

Kelley JL, Morrell LJ, Inskip C, Krause J, Croft DP, 2011. Predation risk shapes social networks in fission-fusion populations. PloS one 6(8): e24280.

Kelley JL, Magurran AE, 2003. Learned predator recognition and antipredator responses in fishes. Fish Fish 4(3): 216–226.

Kimbell HS, Morrell LJ, 2015. Turbidity influences individual and group level responses to predation in guppies, Poecilia reticulata. Anim Behav 103: 179–85.

Krause J, Ruxton GD, 2002. Living in groups. Oxford, UK: Oxford University Press.

Krause J, 1994. The influence of food competition and predation risk on sizeLassortative shoaling in juvenile chub (Leuciscus cephalus). Ethol 96: 105–116.

Kroeker KJ, Sanford E, Jellison BM, Gaylord B, 2014. Predicting the effects of ocean acidification on predator-prey interactions: a conceptual framework based on coastal molluscs. Biol Bull 226 (3): 211–222.

Laidre KL, Stirling I, Lowry LF, Wiig Ø, Heide-Jørgensen MP, Ferguson SH, 2008. Quantifying the sensitivity of Arctic marine mammals to climateLinduced habitat change. Ecol Appl 18(sp2): 97–125.

Lehtonen J, Jaatinen K, 2016. Safety in numbers: the dilution effect and other drivers of group life in the face of danger. Behav Ecol Sociobiol 70: 449–458.

Malavasi S, Cipolato G, Cioni C, Torricelli P, Alleva E, Manciocco A, Toni M, 2013. Effects of Temperature on the Antipredator Behaviour and on the Cholinergic Expression in the European Sea Bass (Dicentrarchus labrax L.) Juveniles. Ethol 119(7): 592–604.

Martin CW, Fodrie FJ, Heck KL, Mattila J, 2010. Differential habitat use and antipredator response of juvenile roach (Rutilus rutilus) to olfactory and visual cues from multiple predators. Oecologia 162(4): 893–902.

Magurran AE, Seghers BH, 1990. Population differences in the schooling behaviour of newborn guppies, Poecilia reticulata. Ethol 84: 334–342.

Magurran AE, Pitcher TJ, 1987. Provenance, shoal size and the sociobiology of predator-evasion behaviour in minnow shoals. Proc R Soc B 229: 439–465.

Mathis A, Smith RJ, 1993. Fathead minnows, Pimephales promelas, learn to recognize northern pike, Esox lucius, as predators on the basis of chemical stimuli from minnows in the pike’s diet. Anim Behav 146(4): 645–456.

McCormick MI, Manassa R, 2008. Predation risk assessment by olfactory and visual cues in a coral reef fish. Coral Reefs 27: 105–113.

Michael SC, Patman J, Lutnesky MM, 2021. Water clarity affects collective behavior in two cyprinid fishes. Behav Ecol Sociobiol 75: 1–13.

Miller N, Gerlai R, 2012. From schooling to shoaling: patterns of collective motion in zebrafish (Danio rerio). PloS one 7: e48865.

Meuthen D, Baldauf SA, Bakker T, Thünken T, 2016. Predator-induced neophobia in juvenile cichlids. Oecologia 181(4): 947–58.

Nafus MG, Germano JM, Swaisgood RR, 2017. Cues from a common predator cause survivallinked behavioral adjustments in Mojave Desert tortoises (Gopherus agassizii). Behav Ecol Sociobiol 71(10): 1–10.

Ofstad EG, Herfindal I, Solberg EJ, Sæther BE, 2016. Home ranges, habitat and body mass: simple correlates of home range size in ungulates. Proc R Soc B 283: 20161234.

Oliveira TA, Idalencio R, Kalichak F, dos Santos Rosa, JG, Koakoski G, et al., 2017. Stress responses to conspecific visual cues of predation risk in zebrafish. PeerJ 5: e3739.

Parra GJ, Corkeron PJ, Arnold P, 2011. Grouping and fission–fusion dynamics in Australian snubfin and Indo-Pacific humpback dolphins. Anim Behav 82(6): 1423–1433.

Pearson HC, 2009. Influences on dusky dolphin (Lagenorhynchus obscurus) fission-fusion dynamics in Admiralty Bay, New Zealand. Behav Ecol Sociobiol 63(10):1437–1446.

Pérez-Escudero A, Vicente-Page J, Hinz RC, Arganda S, De Polavieja GG, 2014. idTracker: tracking individuals in a group by automatic identification of unmarked animals. Nat meth (7): 743–748.

Phen C, Thang TB, Baran E, Vann LS, 2005. Biological reviews of important Cambodian fish species, based on FishBase 2004. World Fish Center, Phnom Penh.

Plaut I, 2001. Critical swimming speed: its ecological relevance. Comp Biochem Physiol Part A Mol Integr Physiol 131(1): 41–50.

Ratcliff R, 1993. Methods for dealing with reaction time outliers. Psychological bulletin 114: 510.

Rieucau G, Fernö A, Ioannou CC, Handegard NO, 2015. Towards of a firmer explanation of large shoal formation, maintenance and collective reactions in marine fish. Rev Fish Biol Fish 25(1): 21–37.

Rifkin JL, Nunn CL, Garamszegi LZ, 2012. Do animals living in larger groups experience greater parasitism? A meta-analysis. Am Nat 180: 70–82.

Riehl R, Kyzar E, Allain A, Green J, Hook M, Monnig L, Rhymes K, Roth A, Pham M, Razavi R, DiLeo J, 2011. Behavioral and physiological effects of acute ketamine exposure in adult zebrafish. Neurotoxicol Teratol 33(6): 658–667.

Rubenstein DI, 1978. On predation, competition, and the advantages of group living. In Social Behaviour, (pp. 205–231). Springer, Boston, MA.

Séguret A, Collignon B, Halloy J, 2016. Strain differences in the collective behaviour of zebrafish Danio rerio) in heterogeneous environment. R Soc Open Sci 3: 160451.

Seigel AR, DeVriendt IG, Strand MC, Shastri A, Wisenden BD, 2022. Association of predation risk ith a heterospecific vocalization by an anabantoid fish. J Fish Biol 100(2): 543–8.

Sekhar MA, Singh R, Bhat A, Jain M, 2019. Feeding in murky waters: acclimatization and landmarks improve foraging efficiency of zebrafish (Danio rerio) in turbid waters. Biol Lett 15: 20190289.

Smith ME, Belk MC, 2001. Risk assessment in western mosquitofish (Gambusia affinis): do multiple cues have additive effects? Behav Ecol Sociobiol 51(1): 101–107.

Snekser JL, Ruhl N, Bauer K, McRobert SP, 2010. The influence of sex and phenotype on shoaling decisions in zebrafish. Int J Comp Psychol 23(1): 70–81.

Speedie N, Gerlai R, 2008. Alarm substance induced behavioral responses in zebrafish (Danio rerio). Behav Brain Res 188: 168–177.

Spence R, Gerlach G, Lawrence C, Smith C, 2008. The behaviour and ecology of the zebrafish, Danio rerio. Biol Rev 83: 13–34.

Stratmann A, Taborsky B, 2014. Antipredator defences of young are independently determined by genetic inheritance, maternal effects and own early experience in mouthbrooding cichlids. Funct Ecol 28: 944–953.

Suriyampola PS, Shelton DS, Shukla R, Roy T, Bhat A, Martins EP, 2016. Zebrafish social behavior in the wild. Zebrafish 13: 1–8.

Szabo B, Mangione R, Rath M, Pašukonis A, Reber SA, Oh J, Ringler M, Ringler E, 2021. Naive poison frog tadpoles use bi-modal cues to avoid insect predators but not heterospecific predatory tadpoles. J Exp Biol 224(24): jeb243647.

Utne-Palm AC, 2002. Visual feeding of fish in a turbid environment: physical and behavioural aspects. Mar Freshw Behav Physiol 35: 111–128.

Videler JJ, Wardle CS, 1991. Fish swimming stride by stride: speed limits and endurance. Rev Fish Biol Fish 1(1): 23–40.

Ward AJ, Mehner T, 2010. Multimodal mixed messages: the use of multiple cues allows greater accuracy in social recognition and predator detection decisions in the mosquitofish, Gambusia holbrooki. Behav Ecol 21(6): 1315–1320.

