## Supplementary file for "Use of multimodal sensory cues in predator avoidance by wild-caught zebrafish shoals"

* Correspondence Address:

ORCID:

Anuradha Bhat: 0000-0002-7447-2380

Ishani Mukherjee: 0000-0002-2279-4169

1. **Calculation justifying why 6.5l of predator cues were added:**

The total volume of the arena was 28l. Therefore by adding 6.5 litres of predator cues, the resultant concentration of cue in the water was around 23% (v/v). Alarm substances, comprised of predator skin and excreta, have indication of the diet of the *Channa* and these odour cues warn prey species of their predator. In a natural stagnant environment, such cues emanating from a predator decreases with distance from a predator and further, due gradual diffusion of water, cue concentration in the water is never 100%. *Channa* individuals, found in pond beds, strike at their prey to catch them and otherwise show little or no movement. Therefore, residing in one place for a long time, it is likely that the zone around them will have odour emanating from them.We assume that the entire arena is the zone in which has the predator odour and, owing to the above factors, we qualitatively set the the cue concentration to around 23%. We maintain this concentration and potency (by controlling *Channa* diet) across all VOC and OC trials.

2. **Control experiments performed showed that gentle addition of water into the arena center had no impact on shoaling properties:**

*Experiment to check whether addition of water impacts shoal properties*

*Experimental set up*

Shoal responses were recorded under two treatments- (1) Controls or no cue treatments (NC) in which shoals were video recorded after placing them in the arena; (2) Water addition treatment (WC) received 6.5l of water - added very gently into the center of the arena. As in OC, VC and VOC, the shoals were video recorded after addition of the cue.

As in the other treatments, the shoal was allowed to acclimatize for 20 minutes following which a water was added or not added, depending on the treatment regime. Shoals that would receive water were put in the arena with water depth ~4cm as the final water depth would become 5cm after water addition. Two minutes after addition of water, (as in other treatments), or immediately after acclimatization in case of NC, the shoal was video recorded for 20 minutes, using a camera (Canon Legria HF R306) placed vertically overhead. We tested 20 shoals, 10 per treatment and each shoal was tested once. The order of treatment-type was random and all analyses was performed by the observer blind to the treatment.

*Analysis*

Inter-individual distances were calculated between shoal members every 10sec. Using the software ImageJ, we obtained the coordinates of the entire shoal (10 fishes) and from these coordinates (following Michael et al. 2021), we calculated the interindividual distances between all members. The mean inter-individual distance for a given shoal was the average of inter-individual for 30 frames.We compared inter-individual distances between treatments (WC and NC) using Wilcoxon Unpaired test and the p -cut off was set at 0.05.

*Results*

The average interindividual distance was comparable between shoals that received no cue and shoals that received water cues (Wilcoxon Unpaired test results: W=48, n=10, p=0.91) (Figure S1).

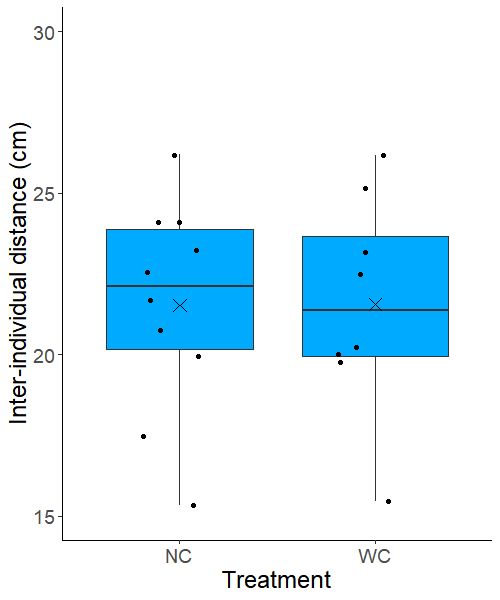

a	a

**Figure S1**: The mean interindividual distances (in cm) are depicted using box-and-whisker plots in which each box represents the interquartile range, the line within the box indicates the median while the whiskers represent the range of the data. Each data point is a dot and the mean is the cross. NC= treatments with no cues; WC= treatments with water cue. Comparisons were performed using Wilcoxon’s Unpaired test (N_NC_ =N_WC_=10, p>0.05).

3. **Experiments to check whether fishes perceive toy predator differently from a real predator**

*Experimental set up*

A 30 × 30 × 50cm tank was taken and divided into two compartments with the help of a transparent partition (i.e. only visual cue exchange was allowed). The tank was filled with aged water up to a height of 10cm. In the smaller compartment (dimensions= 12 × 30 × 30 cm), a toy predator or a real predator was placed. In the larger compartment (dimensions= 38 × 30 × 30cm), an inspection zone was marked 2.5cm (1 body length) next to the smaller compartment and a test fish was released (Fig. S1A). We defined inspection time as the time when the test fish was in the inspection zone. The latency to inspect the predator (toy or real) was the time taken by the test fish to first enter the inspection zone. 14 test fishes (7 for each condition) were exposed to a toy predator or a real predator. Trials were carried out in random order and the observer was blind to the treatment type.

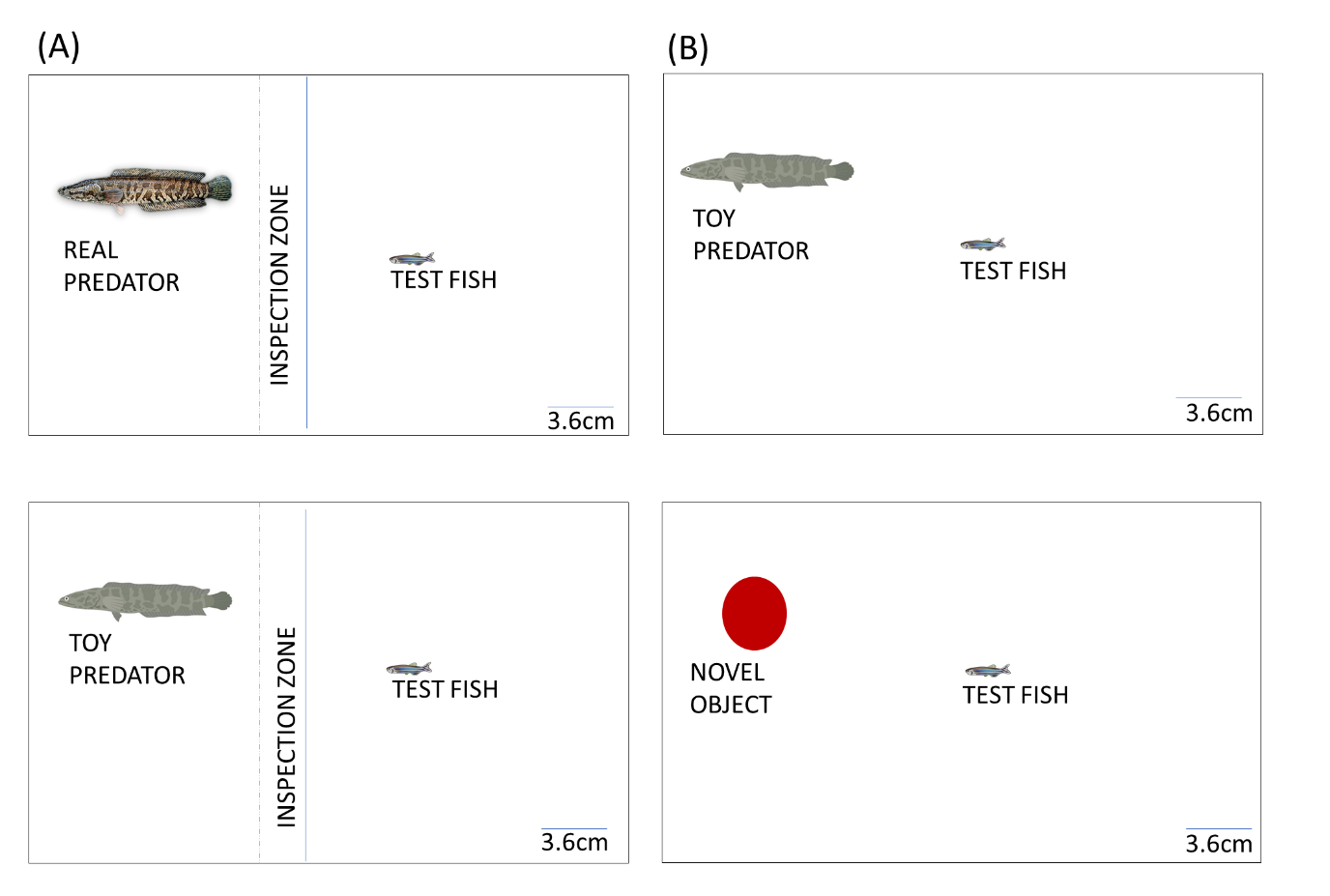

**Figure S2A:** **An overhead view of the arena used to test whether fishes perceive toy predator differently from a real predator.** This schematic has been drawn to scale.

*Results*

In terms of latency to inspect, test fishes showed no difference between a real predator (Mean ±SE=15.87 ± 8.48) and a toy predator (Mean ±SE=24.42±17.16) (Wilcoxon Unpaired test W= 25.5, p= 0.94) (Figure S2B). In terms of number of inspections, test fishes showed no difference between a real predator (Mean ±SE=6.14 ± 1.29) and a toy predator (Mean ±SE=7±1.15) (Wilcoxon Unpaired test W = 20, p= 0.61) (Figure S2C).Thus, from the above experiments we conclude that the behaviour of fishes towards a toy predator is similar to that towards a real predator.

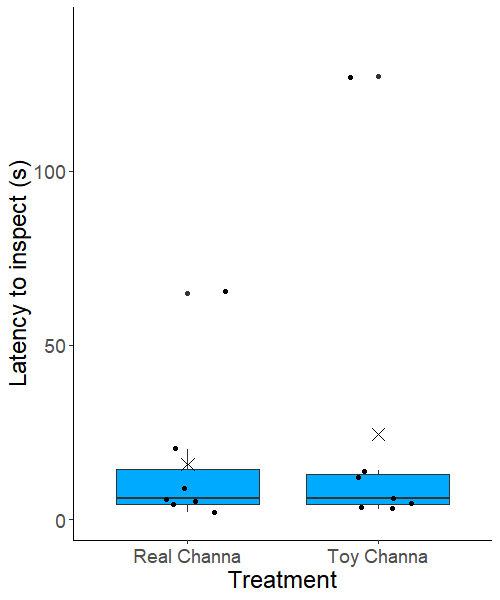

a	a

**Figure S2B:** The latency to inspect (in seconds) is depicted using box-and-whisker plots in which each box represents the interquartile range, the line within the box indicates the median while the whiskers represent the range of the data. Comparisons were performed using Wilcoxon’s Unpaired test (real predator, toy predator=7, p>0.05).

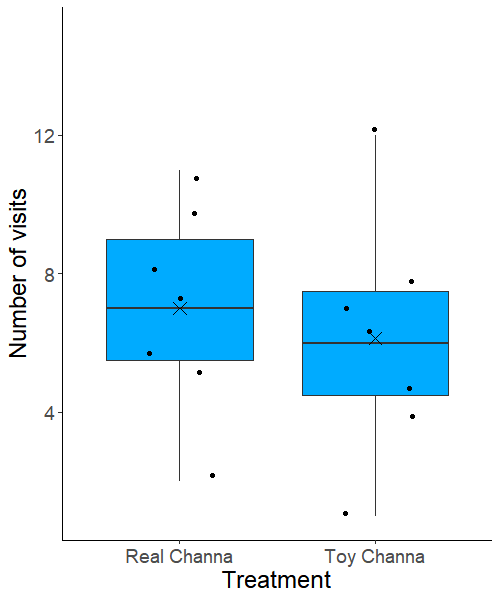

a	a

**Figure S2C:** The number of inspections is depicted using box-and-whisker plots in which each box represents the interquartile range, the line within the box indicates the median while the whiskers represent the range of the data. Comparisons were performed using Wilcoxon’s Unpaired test (real predator, toy predator=7, p>0.05).

4. **Experiments to check whether fishes perceive toy predator differently from a novel object**

*Experimental set up*

A 30 × 30 × 50cm tank was taken and filled with aged water up to a height of 10cm. A toy predator or a novel object was placed at one end of the rectangular tank. An inspection zone was marked 4cm (with pencil on the glass) adjacent to the toy predator or novel object. We defined inspection time as the time when the test fish was in the inspection zone. The latency to inspect the toy predator or novel object was the time taken by the test fish to first enter the inspection zone.

16 test fishes (8 for each condition) were exposed to a toy predator or a novel object. Trials were carried out in random order and the observer was blind to the treatment type.

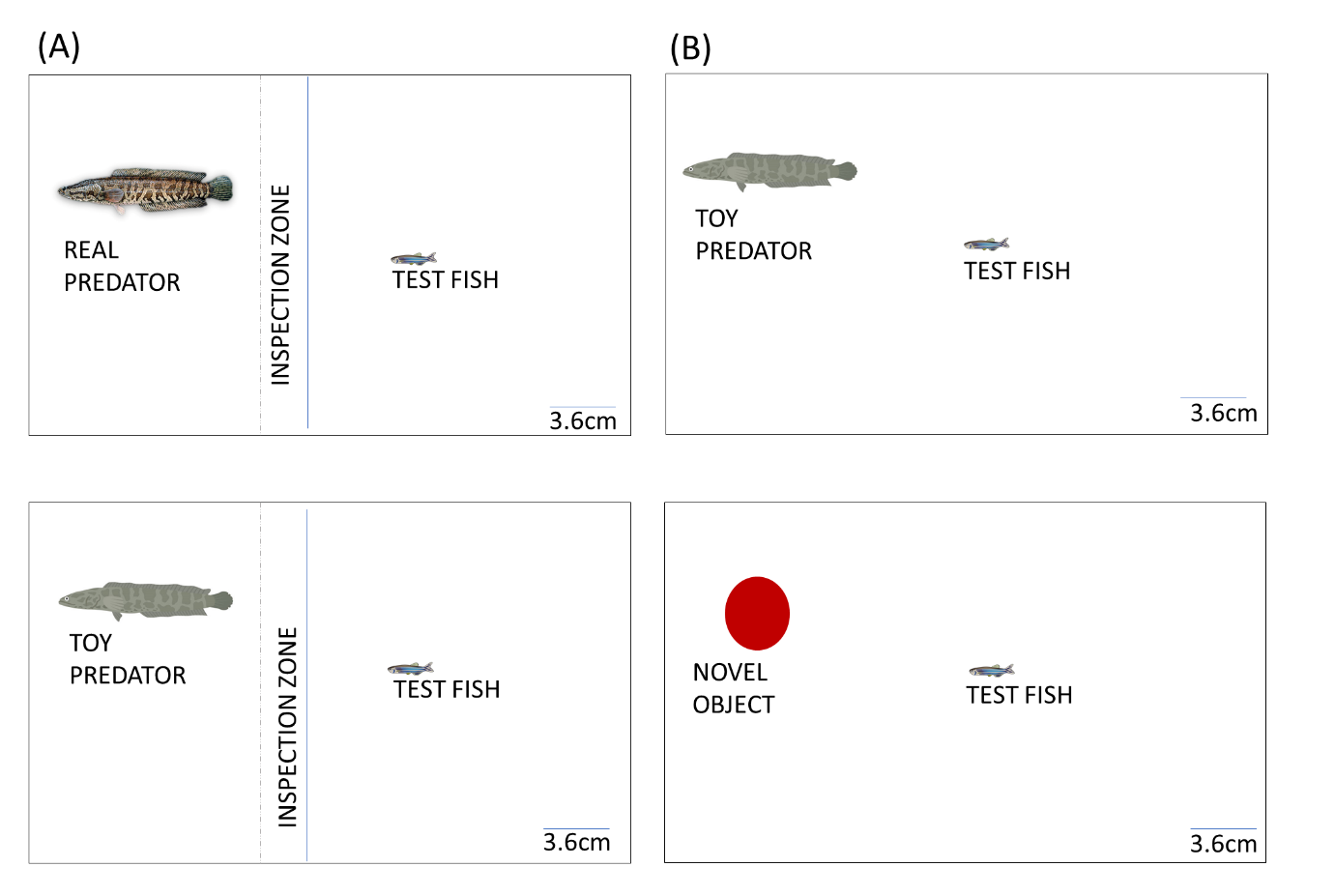

**Figure S3A:** **An overhead view of the arena used to test whether fishes perceive toy predator differently from a novel object.** This schematic has been drawn to scale.

*Results*

In terms of latency to inspect, test fishes visited the toy predator significantly quicker (Mean ±SE=4.75±1.69) than a novel object (Mean ±SE=18±9.68) (Wilcoxon Unpaired test W = 54, p= 0.02) (Figure S3B). In terms of number of inspections, test fishes visited the toy predator (Mean ±SE=7.12±.1.4) comparably as a novel object (Mean ±SE=5.5±0.96) (Wilcoxon Unpaired test W=44, p= 0.22) (Figure S3C).

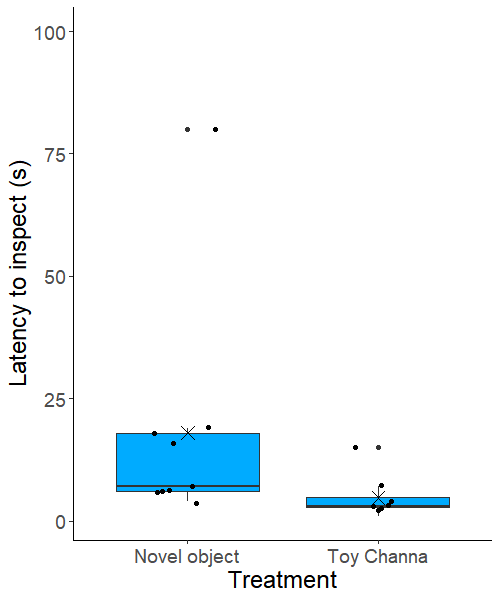

a	b

**Figure S3B:** The latency is depicted using box-and-whisker plots in which each box represents the interquartile range, the line within the box indicates the median while the whiskers represent the range of the data. Comparisons were performed using Wilcoxon’s Unpaired test (object, toy predator=8, p>0.05).

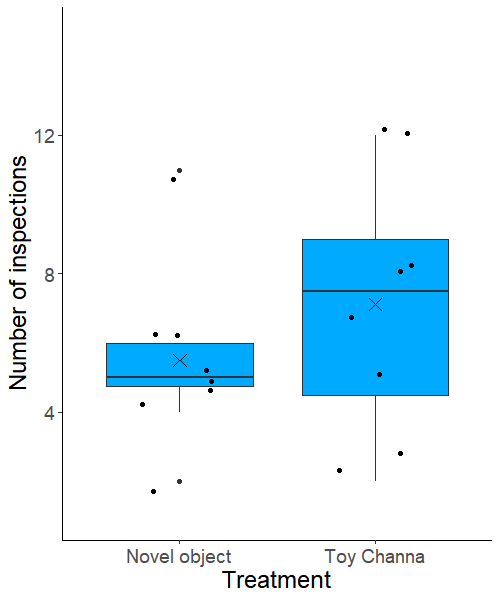

a	a

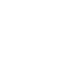

**Figure S3C:** The number of inspections is depicted using box-and-whisker plots in which each box represents the interquartile range, the line within the box indicates the median while the whiskers represent the range of the data. Comparisons were performed using Wilcoxon’s Unpaired test (object, toy predator=8, p>0.05). The box indicates the median while the whiskers represent the range of the data. Comparisons were performed using Wilcoxon’s Unpaired test (object, toy predator=8, p>0.05).

5. **Details of how heat plots showing the distribution of individuals around a focal individual were constructed :**

At a given frame, we obtained the coordinates of all fishes using ImageJ. We chose a given fish (the focal fish F) and we transformed the position of all fishes such that the focal fish is at the origin F(0,0). To do this, we subtracted its coordinate from the coordinates of all other fishes. This gives us the positions of all fishes around the focal fish F(0,0). For all frames, all fishes were considered as focal fishes. We obtained a list of coordinates depicting positions of fishes around a focal fish F. Using a Matlab script, we plotted these coordinates. What we obtain is a visual depiction of inter-individual distances. In our heat maps, using a contour function, we show the proportion of individuals within a given area.

6. **Details of how velocity was plotted:**

Automatic tracking all individuals using idTracker (followed by manual correction using idPlayer) fetched us a trajectory file of each fish. We removed areas not tracked by the software and then divided the data into small snippets such each snippet size would be greater than 5s (125 frames). From that, we have calculated the speed at time t of every fish in each direction (x and y direction) with the help of this general equation of motion:

$$V_{x}\left( t \right)=\frac{x\left( t+\Delta t \right)-x\left( t \right)}{\Delta t}$$

$$V_{y}\left( t \right)=\frac{y\left( t+\Delta t \right)-y\left( t \right)}{\Delta t}$$

here time interval $\Delta t$=0.04 s

After that, we then calculate the magnitude of velocity from snippet V_x_. and V_y_ using this formula:

$$V(t)=\sqrt{V_{x}^{2}(t)+V_{y}^{2}(t)}$$

To smoothen our data, we did a moving average of speed at the level of individual components x and y after equations (1) and (2). Then as the usual way, we calculated the magnitude of the velocity (or speed). The algorithm we used for calculating the moving average 10 frame speed V_ma_ at time point is:-

$$V_{ma}(t_{p})=\sqrt{\left( \frac{1}{10}\sum_{i=t_{p}-9}^{t_{p}} V_{x}(i) \right)^{2}+\left( \frac{1}{10}\sum_{i=t_{p-9}}^{t_{p}} V_{y}(i) \right)^{2}}$$

Here t_p_ = a Time point is an integer number, t_p_≥10.

7. **Our criteria for dividing the velocity into probability of acceleration, deacceleration and freezing phases:**

Acceleration, deacceleration or freezing were calculated in the following way: if the velocity increased for atleast 5 consecutive frames (200ms) we termed this phase as acceleration. If it decreased, we termed it deceleration. Phases were designated as freezing if for 5 consecutive frames, the velocity of the fish was equal to or less than 2.5 cm/s (i.e. 1bodylength/s). Thus, to impose stringency and discard noise, only if the phases were equal to 200ms or more than these were considered this to be either of these three phases. The duration of the acceleration, deceleration and freezing events were calculated. The duration of a given phase divided by the total duration (i.e. the duration of all three phases) fetched us the proportion time spent in the given phase.

8. **Proportion time spent in acceleration and deacceleration phase across the four treatments**

GLM revealed proportion time spent in: (A) acceleration phase was not dependent on the cue provided (GLM: Wald type IIχ^2^ = 3.36, df = 3, p=0.30, Table S1) (B) deacceleration phase was not dependent on the cue provided not on the cue provided (GLM: Wald type IIχ^2^ = 2.27, df = 3, p=0.51, Table S2) (Figure S4).

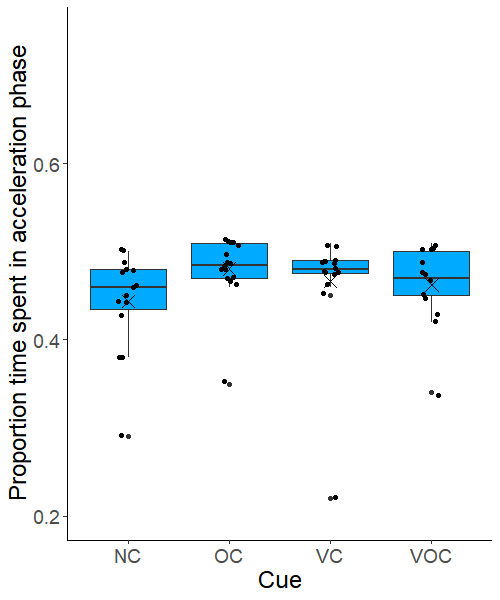

a	a	a	a

Proportion time spent in acceleration phase

**Figure S3B:** The proportion time spent in acceleration phase has been depicted using box-and-whisker plots in which each box represents the interquartile range, the line within the box indicates the median while the whiskers represent the range of the data. Data points have been represented as dots. The letter ‘a’ placed above the boxes represents no significant difference between the categories Comparisons were performed using GLM (Sample size: NC, OC, VC=14 shoals, VOC=13 shoals; p>0.05).

a	a	a	a

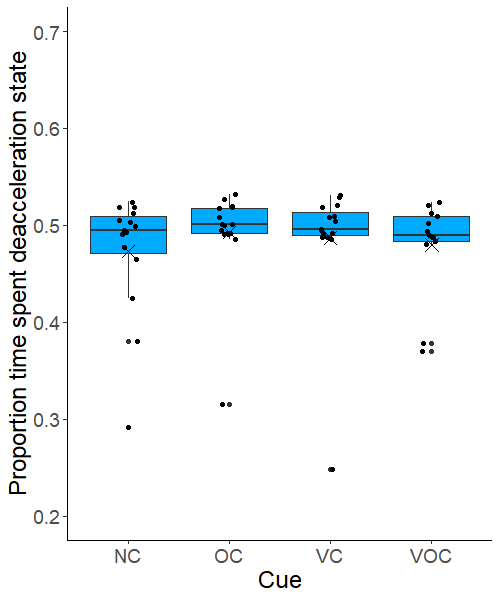

Proportion time spent in de-acceleration phase

**Figure S3B:** **:** The proportion time spent in de-acceleration phase has been depicted using box-and-whisker plots in which each box represents the interquartile range, the line within the box indicates the median while the whiskers represent the range of the data. Data points have been represented as dots. The letter ‘a’ placed above the boxes represents no significant difference between the categories. Comparisons were performed using GLM (Sample size: NC, OC, VC=14 shoals, VOC=13 shoals; p>0.05).
